# Valosin-containing protein regulates the stability of amyotrophic lateral sclerosis-causing fused in sarcoma granules in cells by changing ATP concentrations inside the granules

**DOI:** 10.1101/2021.12.24.474151

**Authors:** Kyota Yasuda, Tomonobu M. Watanabe, Myeong-Gyun Kang, Jeong Kon Seo, Hyun-Woo Rhee, Shin-ichi Tate

## Abstract

Fused in sarcoma (FUS) undergoes liquid-liquid phase separation (LLPS) to form granules in cells, leading to pathogenic aggregations that cause neurodegenerative diseases, including amyotrophic lateral sclerosis (ALS). Proteomics analysis revealed that FUS granules contain valosin-containing protein (VCP), a member of the AAA family ATPase. Confocal microscopy images showed that VCP co-localized in the FUS granules in cells. This study demonstrates that VCP in granules has a two-faced role in FUS granulation: VCP stabilizes *de novo* FUS granules, while VCP present in the granules for extended periods dissolves them. This VCP function relies on its ATPase activity to consume ATP in granules. VCP stabilizes *de novo* FUS by reducing intragranular ATP concentrations to a range below the cytosolic concentration. VCP continually consumes ATP during its stay in the granules, which eventually lowers ATP concentrations to a range that destabilizes the granules. VCP, therefore, acts as a timer to limit the residence of FUS granules in cells and thereby prohibits the FUS fibrillization that occurs in persistent granules. VCP ATPase activity plays a role in FUS granule turnover.

**Summary statement:** VCP recruited to FUS granules regulates the stability of the granules in a time-dependent manner by consuming intragranular ATP with its ATPase activity.

## Introduction

Fused in sarcoma (FUS), also known as translated in liposarcoma, is a ubiquitously expressed DNA/RNA-binding protein that is predominantly localized in the cell nucleus (Crozat et al., 1993; Lee et al., 2006). FUS is involved in DNA repair and transcription regulation, RNA splicing, and RNA transport (Iko et al., 2004; Mackenzie and Neumann, 2017). FUS achieves liquid-liquid phase separation (LLPS) to form liquid droplets with RNA and other proteins (RNP granules) under physiological conditions in the cytoplasm (Patel et al., 2015b). Aberrant RNP granulation is linked to neurodegenerative diseases, including amyotrophic lateral sclerosis (ALS) and frontotemporal lobar degeneration (FTLD) (Chen et al., 2019; Verdile et al., 2019).

ALS-causing mutations are mapped over FUS primarily localized in the C-terminal nuclear localization signal (NLS) with motifs R/H/KX_2-5_PY and PY-NLS (Chen et al., 2019; Kwiatkowski et al., 2009; Lee et al., 2006; Naumann et al., 2019; Shang and Huang, 2016; Vance et al., 2009). FUS variants with mutations in NLS mislocalize to the cytoplasm, causing a liquid-to-solid phase transition to form insoluble pathogenic aggregates (Marrone et al., 2019; Shelkovnikova et al., 2014).

Cytoplasmic mislocalized FUS is also associated with stress granules (SGs) (Daigle et al., 2013; Sama et al., 2013). Cytoplasmic FUS has been isolated as a component in an RNA-transporting granule associated with kinesin-1, suggesting that FUS is involved in regulating motor motility (Kanai et al., 2004; Yasuda et al., 2013). Mislocalized FUS variants were found to co-localize with kinesin-1 and mRNA in SGs, suggesting that SG formation arrests RNA transport for dysregulating RNA translation in axons (Yasuda et al., 2017).

Mutations that promote FUS granulation cause neurodegeneration (Jackrel and Shorter, 2014; López-Erauskin et al., 2018; Murakami et al., 2015; Ryan et al., 2019; Sun et al., 2011). Therefore, FUS mutants with elevated granulation propensity promote ALS pathogenesis. Isolated FUS undergoes LLPS *in vitro* (Chong et al., 2018; Kato et al., 2012). However, the regulatory mechanism of FUS granulation, in which various co-localizing components in FUS granules engage, remains elusive (Calabretta and Richard, 2015).

Valosin-containing protein (VCP) is a genetic marker of neurodegenerative diseases, including ALS (Johnson et al., 2010; Watts et al., 2007). VCP works in multiple cellular processes (Stolz et al., 2011; Sun and Qiu, 2020). It plays a central role in the ubiquitin-dependent proteasome pathway (Dai et al., 1998; Dai and Li, 2001; Sun and Qiu, 2020; Xia et al., 2016). VCP is a proteolysis-promoting factor that plays a role in clearing SGs in cells (Buchan et al., 2013). VCP localizes FUS in the nucleus; the A232E mutant VCP prompts cytosolic mislocalization of FUS (Tyzack et al., 2019). VCP may drive clathrin-dependent endocytosis; VCP-knockdown cells show impaired endocytosis (Liu et al., 2020a). Soluble FUS are supposed to be cleared by endocytosis, where VCP promotes their clearance by activating the endocytic pathway (Ju et al., 2009; Liu et al., 2020a; Wang et al., 2015). Sequestrating VCP in FUS aggregates thus impairs endocytosis to reduce FUS clearance from the cytosol (Liu et al., 2020a).

VCP belongs to the type II class of the AAA (ATPase associated with various activities) ATPase family (Xia et al., 2016). VCP is a homohexamer protein, and each subunit has two AAA ATPase domains (Xia et al., 2016). VCP rich in ATPases co-localized to FUS granules should consume substantial amounts of ATP in the granules.

In this study, we explored the role of VCP co-localized in FUS granules in cells. This study illustrates that VCP in *de novo* FUS granules rigidifies the granules. However, the extended residence of VCP in the granules destabilized the granules. VCP works as a timer to limit the lifetime of FUS granules in cells. This study adds a new role of VCP in protein granulation in cells.

## Results

### Fused in sarcoma granules contain valosin-containing protein in cells

To map the interactome of FUS, we introduced spatiotemporal proximity cross-linking *via* light activation. This method utilizes a photoreactive aryl azide-conjugated ligand (PL) covalently bound to HaloTag. The aryl azide moiety of the PL at the bait protein surface is converted to reactive nitrene when UV illuminated. The nitrene chemically links proteins in spatial proximity (within a few Å) to the HaloTag-conjugated bait protein (Mishra et al., 2022). Halo-FLAG-FUS complexed with PL in NIH/3T3 cells captures physically interacting proteins of FUS upon UV illumination (Fig. S1A).

Halo-FLAG-FUS cross-linked proteins after UV light illumination, as observed in the western blot (Fig. S1B). The proteins cross-linked to Halo-FLAG-FUS were collected by immunoprecipitation using an anti-FLAG antibody. The collected proteins were subjected to mass analysis after trypsin-digestion. Fold changes in eight representative proteins after UV illumination are compared (Fig. S1C).

We also conducted proximity cross-linking experiments on Halo-FLAG-FUS P525L, a granulation-prone mutant FUS (Fig. S1C). Among the proteins cross-linked to FUS, VCP showed a significant reduction in affinity to FUS P525L (Fig. S1C). Immunoprecipitation with an anti-FLAG antibody to the lysate from NIH/3T3 overexpressing the FLAG-FUS P525L showed a reduced co-precipitated VCP relative to the wild-type FUS (Fig. 1A), confirming the reduced VCP affinity to FUS P525L found in the proximity cross-linking experiments (Fig. S1C).

**Figure 1:**
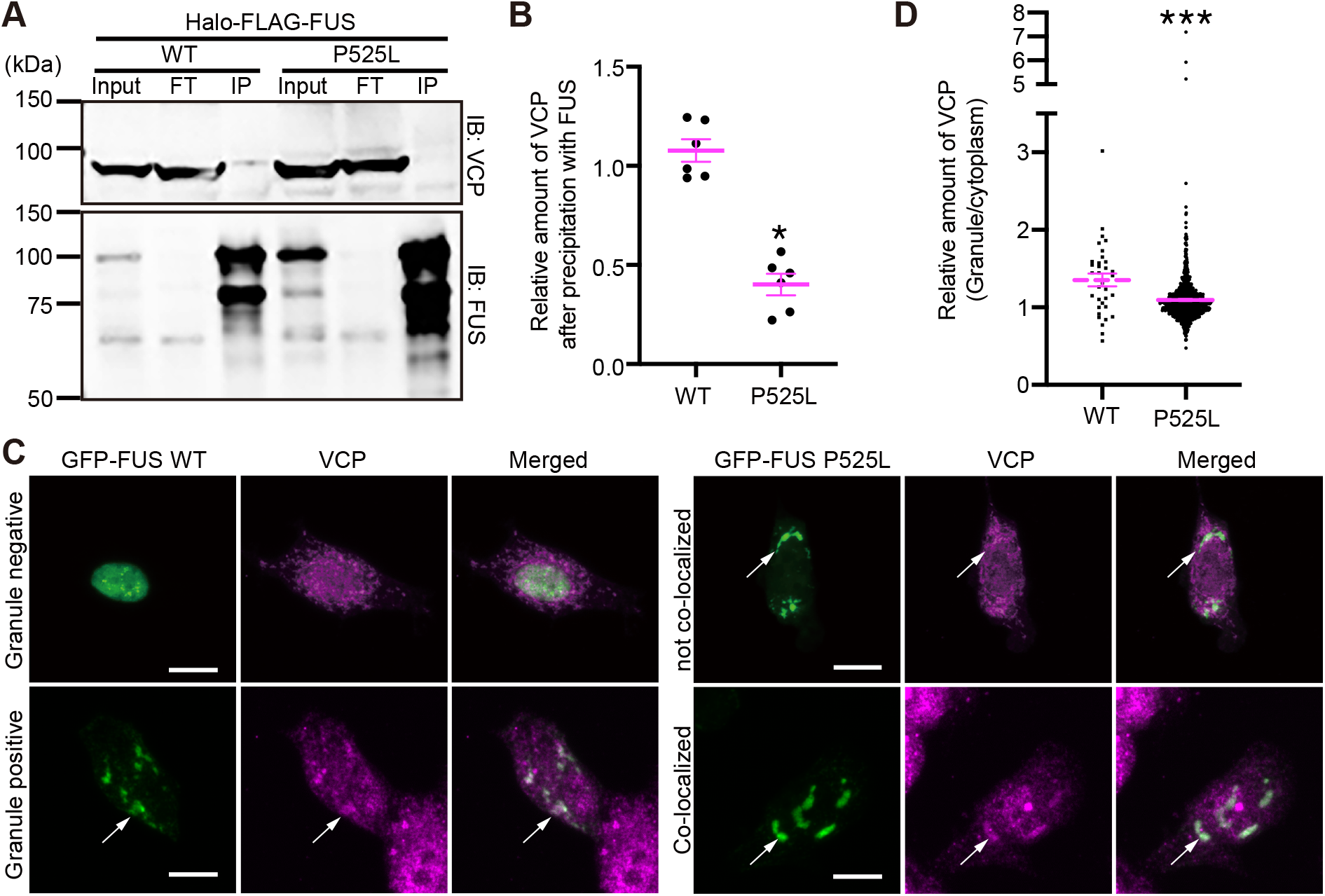
ALS-related mutation on FUS reduces the interaction between FUS and VCP. (A) Western blotting for immunoprecipitated samples of FLAG-tagged FUS protein overexpressed in cells. The lysate without precipitation (input), supernatant after precipitation (FT), or precipitates (IP) were loaded and detected with anti-VCP antibody (upper) or anti-FUS antibody (lower). The observed bands around 100 kDa in the FUS-immunoreacted membrane are the Halo-FLAG-FUS, and the band between 75 kDa and 50 kDa is the endogenous FUS. The bands at approximately 75 kDa were non-specific. (B) Relative amount of VCP precipitated with Halo-FLAG-FUS. The intensities of VCP-bands were divided by the precipitated Halo-FLAG-FUS intensity and normalized to the average of the WT samples. \**P* = 0.0312 by Wilcoxon matched-pairs signed rank test, mean*±*s.e.m., six independent experiments. (C) Representative images of immunofluorescence with anti-VCP antibody in cells expressing wild-type (GFP-FUS WT) or mutant (GFP-FUS P525L) FUS. Upper images of GFP-FUS WT are of the cell without granules, and the bottom images of GFP-FUS WT are of the cell with granules. Upper images of GFP-FUS P525L are of the cell without co-localization of VCP in granules. The bottom images of GFP-FUS P525L are of the cell with co-localization of VCP in granules. Arrows point to one of the FUS granules. The bars on the images are 10 μm. (D) Quantitative analysis of VCP localization in FUS granules. The ratio of the mean intensity of VCP-immunostaining in granules to the adjacent cytoplasm was calculated. ***P < 0.0001 by Mann-Whitney test, mean*±*s.e.m., n = 966 for P525L, n = 34 for WT samples, two independent experiments. Because the wild-type FUS is less likely to form granules, the n number of wild-type FUS was much smaller than that of mutant FUS.

The cells expressing wild-type FUS containing cytoplasmic granules populate about 15% (Fig. S2A). The population of the granule-containing cells increased to approximately 75% in the cells expressing the FUS P525L (Fig. S2A), as previously reported (Yasuda et al., 2017; Yasuda et al., 2013).

Immunofluorescent microscopic images demonstrated that the endogenous VCP distributed throughout the regions of the cells overexpressing either the wild-type or FUS P525L (Fig. 1C). The fluorescence intensity ratios of the VCP co-localized in the FUS granules against the VCP out of the granules showed that VCP tends to co-localize to the wild-type FUS granules over the FUS P525L mutant granules (Fig. 1D). The co-precipitated VCP with FUS P525L was half of that with the wild-type in amount (Fig. 1B). Immunofluorescence analysis, however, showed VCP co-localized to the FUS P525L granules was reduced by approximately 20% from that of the wild-type (Fig. 1D). The smaller change in the co-localized VCP in FUS P525L granules (Fig. 1D), although VCP has a 50% reduced affinity to the mutant FUS relative to the wild-type (Fig. 1B), suggests that VCP recruitment relies not solely on the binding to FUS but also on the interaction with the other components in the FUS granules.

FUS granulation does not require the presence of VCP. A partial siRNA knockdown of VCP in NIH/3T3 cells (Fig. S3A), where the VCP mediates ubiquitination at a comparable level in normal cells (Dai et al., 1998; Dai and Li, 2001) (Fig. S3B), does not change the population of granule-containing cells (Fig. S3C). It should be noted here that the excessive VCP knockdown severely reduced cell viability. We, therefore, applied the partial VCP knockdown.

### Valosin-containing protein reduces fused in sarcoma dynamics in the granules

VCP overexpression in cells lowers FUS exchanging between inside and out of the granules, as revealed by fluorescence recovery after photobleaching (FRAP) (Axelrod et al., 1976). We used cells overexpressing Green Fluorescent Protein (GFP)-tagged FUS P525L (GFP-FUS P525L) and Red Fluorescent Protein (RFP)-tagged VCP (RFP-VCP). Confocal images of the cells showed that a fraction of RFP-VCP co-localized to the GFP-FUS P525L granules (Fig. 2A).

**Figure 2:**
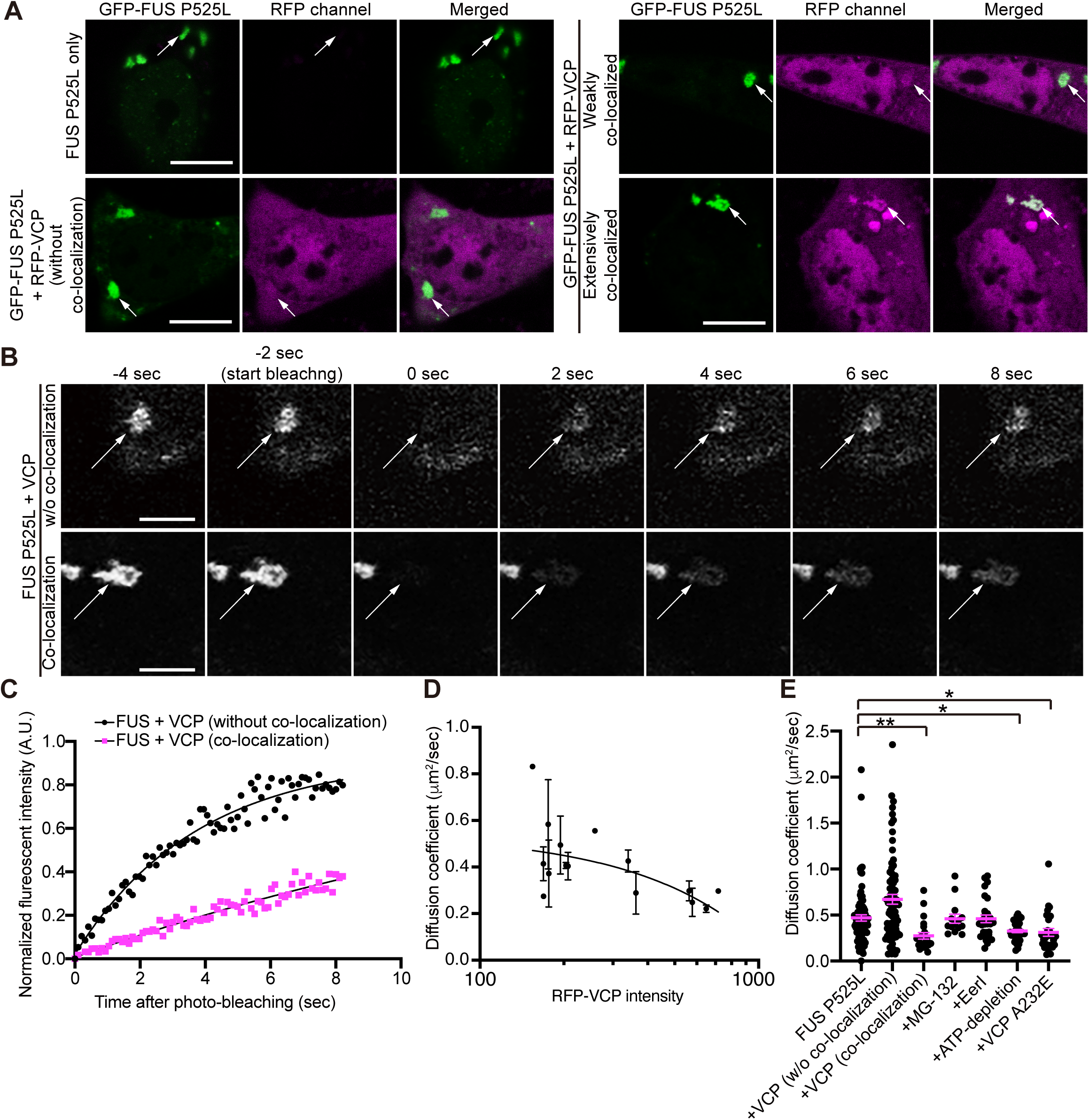
Overexpression of VCP or ATP depletion affects the mobility of FUS granules. (A) Representative images of cells expressing GFP-FUS with (left bottom and right panels) and without RFP-VCP (Left top) co-expression. We showed different levels of co-localization of VCP in the FUS granule, such as without co-localization (left bottom), weak co-localization (right top), and extensive co-localization (right bottom). Arrows indicate one of the FUS granules, for example. (B) Representative images of granules upon FRAP experiments. The granules are from the same cells in (A). The time indicated above is before and after photo-bleaching centered just after bleaching as zero. Arrows indicate the granules targeted for FRAP. (C) The representative photo-recovery graph was generated from the granules data shown in (B). The intensity in the FRAP area after photo-bleaching was normalized and plotted over time to show the example of photo recovery. The fitting curve is also described (line among plots). (D) The data from one trial of FRAP used in Figure 2E (+VCP without co-localization) were used for plotting the diffusion coefficient against exogenous VCP intensity. Each data point is composed of the FRAP data in individual cells (e.g., data point without error bar means that we acquired FRAP data from a single granule in the cell). The black line is the linear regression line. Error bars show s.e.m. Bars on images indicate 10 μm (A) or 5 μm (B). Error bars show s.e.m. (E) The graph of the diffusion coefficient was calculated with FRAP data from cells with below-indicated manipulation. Dunn’s multiple comparison test was performed by comparing FUS P525L to others [**P = 0.0014, *P = 0.0408 (+ATP-depletion), *P = 0.0108 (+VCP A232E), and P > 0.05 for others]. mean*±*s.e.m., n = 86, 76, 22, 14, 28, 39, and 27 from left to right, two independent experiments for +EerI, +MG-132, +ATP-depletion, and +VCP A232E, three independent experiments for others.

In the following experiments, we used FUS P525L because FUS P525L forms a greater number of granules in cells than the wild-type, which allows for the sampling of many data to explore the states of the granules (Fig. S2A). Herein, we refer to FUS P525L as FUS, unless otherwise stated.

The fluorescence recovery rates from GFP-FUS in the granules containing RFP-VCP were lower than those for GFP-FUS without co-localized RFP-VCP (Fig. 2C). The FRAP data demonstrated that VCP in the granules reduced the FUS dynamics. Co-localization of VCP rigidifies the FUS granules.

GFP-FUS diffusion coefficients decreased with increasing RFP-VCP intensities in the granules (Spearman correlation coefficient, *r* = -0.6000, \**P* = 0.0203) (Fig. 2D), suggesting that co-localizing VCP reduces the FUS dynamics in the granules.

### Valosin-containing protein rigidifies fused in sarcoma granules by lowering adenosine triphosphate concentrations inside the granules

VCP is a hexameric protein with two ATPase domains in each subunit (Meyer et al., 2012). VCP performs ATP hydrolysis to segregate ubiquitinated proteins from various cellular complexes to promote their degradation in the ubiquitin-proteasome pathway (Meyer et al., 2012; Stolz et al., 2011).

Two types of proteasome inhibitors, MG-132 and Eeyarestatin I (EerI), were applied to the cells to assess if the proteolysis-promoting function of VCP affects FUS granulation (Fig. 2E). MG-132 blocks the 26S proteasome activity (Jain et al., 2016; Jiang et al., 2013). EerI blocks VCP ATPase activity to inhibit its protein segregation activity (Wang et al., 2008; Wang et al., 2009).

MG-132 marginally changed diffusion coefficients of GFP-FUS in the granules (Fig. 2E), implying that the change in the FUS dynamics in the VCP co-localizing granules does not rely on the VCP-mediated proteolysis process. This is a stark contrast to the reported role of VCP in the SG clearance, where VCP promotes bringing the ubiquitinated proteins to autophagy (Buchan et al., 2013).

Blocking the ATPase activity of VCP by EerI caused no apparent change in the FUS dynamics in the granules (Fig. 2E). VCP unfolds ubiquitinated proteins using ATPase activity to facilitate their proteolysis (Stolz et al., 2011). The results demonstrate that VCP-mediated proteolysis is not linked to FUS dynamics in the granules.

ATP depletion in cells with carbonyl cyanide m-chlorophenyl hydrazone (CCCP) significantly reduced the GFP-FUS diffusion coefficient (Fig. 2E). CCCP blocks the glycolytic pathway to produce ATP, thus causing ATP depletion in cells during culture (Jain et al., 2016).

The diffusion coefficients of GFP-FUS in the ATP depleted cells were close to the values observed for FUS in the VCP co-localized granules (Fig. 2E). These results suggest that VCP ATPases reduce ATP concentrations in the granules by consumption, thereby rigidifying the granules.

Overexpressed VCP distributes uniformly inside cells, and a fraction of them co-localized to the FUS granules (Fig. 2A), as found for the endogenous VCP (Fig. 1C). FUS granules without observable VCP co-localization in the cells overexpressing VCP elevated diffusion coefficients over the FUS granules in the cells without overexpressing VCP (Fig. 2E). VCP in the cytosol may facilitate proteolysis of FUS granules as in the SG clearance by VCP, although its play is less obvious in clearing FUS granules (Buchan et al., 2013). Overexpression exaggerates the proteolysis function of the cytosolic VCP in destabilizing the FUS granules, although clearing FUS granules by the endogenous VCP is not apparent (Fig. 2E).

VCP A232E loses its protein unfolding function but retains three times higher ATPase activity (Gitcho et al., 2009; Halawani et al., 2009; Ju et al., 2009; Meyer and Weihl, 2014; Niwa et al., 2012). Overexpressed VCP A232E reduced the diffusion coefficients of GFP-FUS to the values observed for the FUS granules containing VCP (Fig 2E). The used VCP A232E did not contain RFP for fluorescence detection. The values for the cells overexpressing VCP A232E contain the data from the FUS granules with and without VCP A232E. VCP with higher ATPase activity (Niwa et al., 2012) explains that the FUS dynamics in the cells expressing VCP A232E is close to the value for the cells in ATP depletion (Fig. 2E).

Taken together, VCP in the granules alters FUS dynamics through its ATPase activity to reduce ATP concentrations but not through its proteolysis-promoting function (Fig. 2E).

ATP concentrations change the stability of FUS granules: ATP at low concentrations facilitates LLPS of FUS, while it dissolves the FUS granules at high concentrations (Kang et al., 2019a; Kang et al., 2018; Patel et al., 2017). Kang et al. reported that FUS granulation starts at 0.2 mM ATP concentrations, while the dissolution of the granules starts at 4.0 mM (Kang et al., 2018). They also showed that the FUS granules had maximum sizes at 1.0 mM ATP concentrations, implying that the FUS granules are stable at ATP concentrations of approximately 1.0 mM (Kang et al., 2018).

ATP concentrations in the cytosol of cultured cells range from 3.7 mM–4.1 mM (Yoshida et al., 2016). Therefore, *de novo* FUS granules in cells may retain high ATP concentrations destabilizing the granules. The co-localized VCP consumes ATP in the FUS granules to stabilize the granules by reducing inside ATP concentrations.

### Valosin-containing protein reduces adenosine triphosphate concentrations in cells

The change in ATP concentration by VCP was explored using the Förster resonance energy transfer (FRET)-based indicator, ATeam (Imamura et al., 2009). ATeam changes its emitting fluorescence according to the ATP concentration. In the absence of ATP, ATeam emits 475 nm Cyan Fluorescent Protein (CFP) fluorescence, and it emits 527 nm fluorescence from Yellow Fluorescent Protein (YFP) excited through FRET from CFP in the presence of ATP (Imamura et al., 2001). The fluorescence intensity ratio between 457 nm and 527 nm changes in proportion to the ATP concentration (Imamura et al., 2009): the fluorescence intensity ratio between CFP and YFP, I(CFP)/I(YFP), becomes smaller at higher ATP concentrations.

ATP concentrations monitored by ATeam in cells were compared among the cells expressing RFP-FUS alone and RFP-FUS + VCP and cells in ATP depletion (Fig. 3A). I(CFP)/I(YFP) values for the cells in ATP depletion significantly diminished (Fig. 3B). The cells expressing RFP-FUS alone showed a marginal change in ATP concentration relative to that in cells harboring only ATeam (Fig. 3B). It is noted that ATeam does not co-localize to the FUS granules and thus cannot measure the ATP concentrations inside the granules (Fig. 3A); I(CFP)/I(YFP) gives the average ATP concentrations in the cytosol (Fig. 3B).

**Figure 3:**
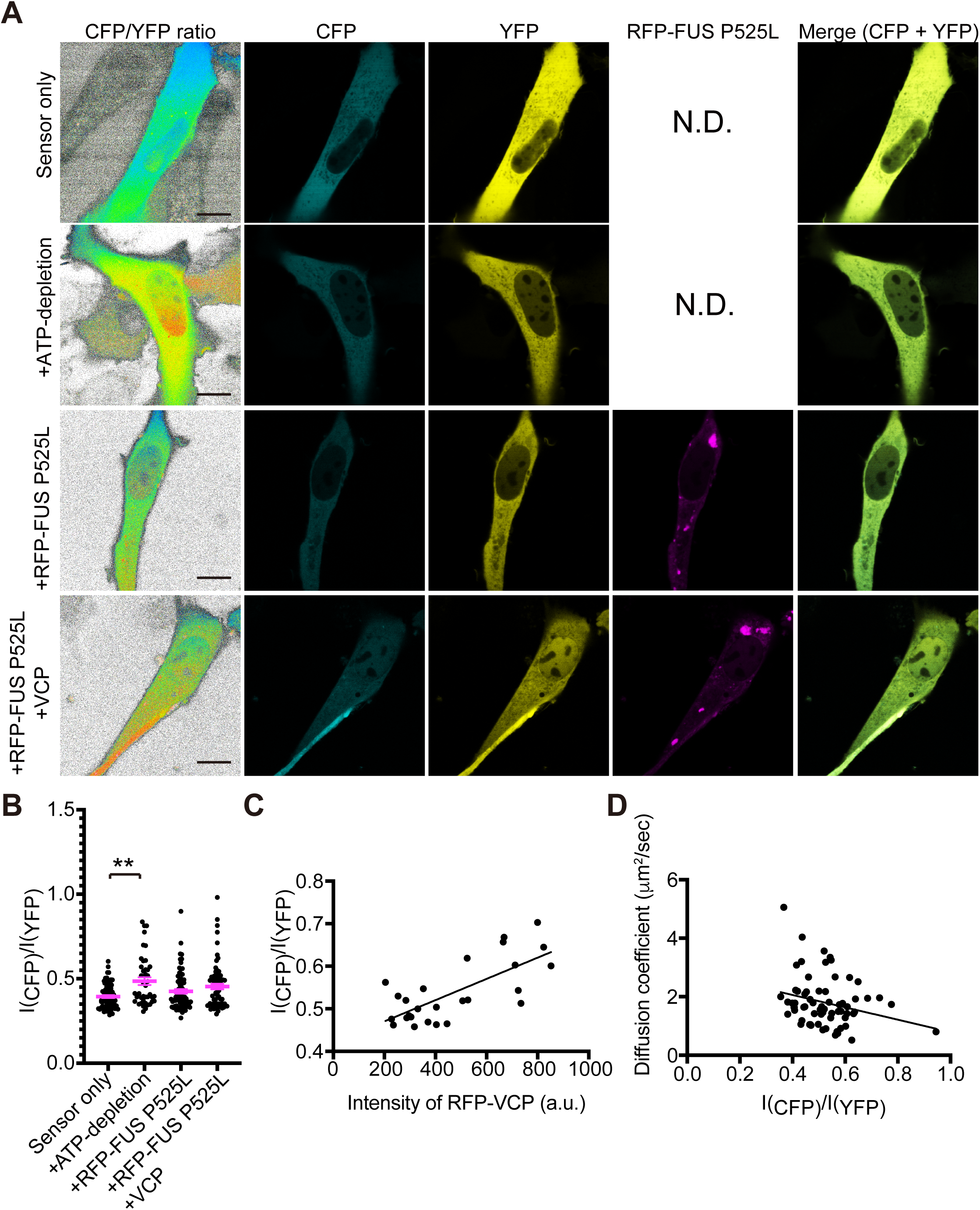
Measurement of ATP concentration with FRET sensor revealed possible relation among VCP amount, ATP concentration in cells, and FUS diffusion coefficient in granules. (A) Representative images of cells with indicated manipulation. ATP-sensor means cells expressing ATeam only. The left panel is the 16 color-image to show the calculated ratio of CFP signals over YFP signals. (B) The FRET-based ATP amount among cells with indicated manipulation. I(CFP) or I(YFP) is the CFP or YFP fluorescence intensity, respectively. **P = 0.0025, or P > 0.05 for others by Dunn’s multiple comparisons test by comparing FUS P525L to others, mean±s.e.m., n = 75, 42, 73, and 69 from left to right. All data were acquired from two individual experiments. (C) The plot of ATeam ratio [I(CFP)/I(YFP)] against RFP-VCP intensity. Each point is from individual cells. N = 26 from one experiment. The black line indicates the linear regression line. (D) The plot of the diffusion coefficient of FUS in granules analyzed by FRAP against ATeam ratio [I(CFP)/I(YFP)] of analyzed FUS granules. Each point is from individual cells. N = 77 from two individual experiments. The black line indicates the linear regression line.

Overexpression of RFP-VCP in cells reduced ATP concentrations in proportion to its expression level (Spearman correlation coefficient, *r* = 0.5740, \*\**P* = 0.0022 (Fig. 3C)).

GFP-FUS diffusion coefficients diminished according to the ATP concentrations in cells (Spearman correlation coefficient, *r* = -0.2391, \*\**P* = 0.0363 (Fig. 3D)). VCP ATPase consumes cytosolic ATP to reduce the ATP concentration in cells to rigidify the FUS granules in proportion to the amount of VCP in cells.

### Valosin-containing protein enlarges the core spheres of fused in sarcoma granules

The granules in the cells are supposed to consist of a liquid-like outer sphere and a solid-like core (Hofweber et al., 2018; Shiina, 2019). Applying digitonin, a detergent, to the cells harboring granules reduces their size by dissolving the liquid-like outer sphere while maintaining the solid-like inner core (Shiina, 2019).

The cells harboring GFP-FUS granules permeabilized with digitonin reduced the granule sizes (Fig. 4A); the liquid-like outer sphere of the FUS granules was dissolved by the detergent (Shiina, 2019). Digitonin reduced granule size by approximately 50% in 20 min after permeabilization (Fig. 4B).

**Figure 4:**
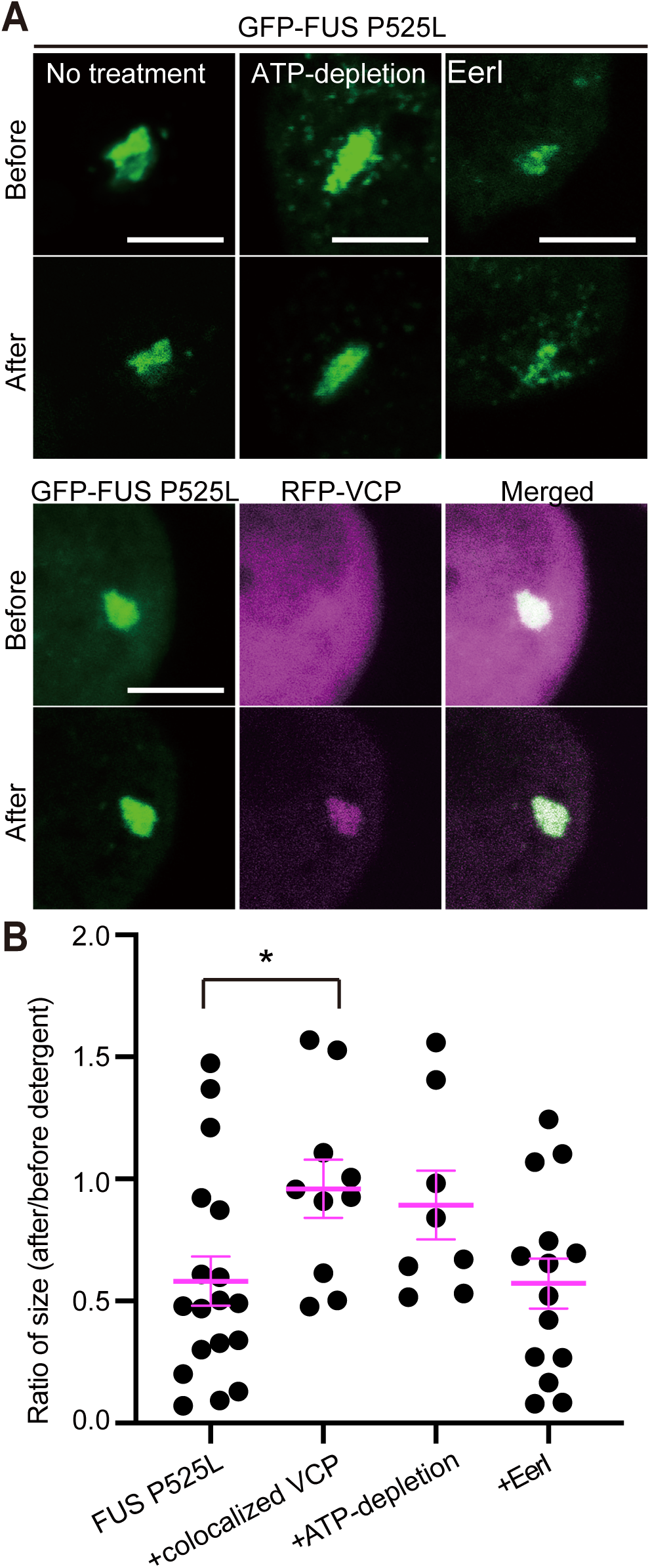
FUS granules retained in cells with overexpression of VCP or ATP-depletion are composed of the less dynamic core sphere. (A) Representative images of FUS granules in cells before and after detergent treatment with and without overexpressing VCP or indicated manipulation. Bars on images show 5 μm. (B) The size changes of FUS granules before and after detergent treatment (20 min). *P = 0.0433, or P > 0.05 for others by Dunn’s multiple comparisons test by comparing FUS P525L to others, mean±s.e.m., n = 18, 10, 8, and 14 from left to right. Error bars show s.e.m.

The cells in ATP depletion showed a limited change in granule size (Fig. 4A). The granules in the cells treated with Eerl, VCP ATPase blocker, reduced granule size (Fig. 4A). The statistical data supported the observations of representative images (Fig. 4B). Lowered cytosolic ATP concentrations stabilized the FUS granules while blocking the proteolysis-promoting function of VCP did not change the stability of the FUS granules.

The FUS granules with co-localized RFP-VCP mostly retained their sizes after digitonin permeabilization (Fig. 4A), which is close to the observation for the cells in ATP depletion (Fig. 4B). The results suggest that VCP in the FUS granules rigidifies the liquid-like outer spheres to make them resistant to dissolution by digitonin by reducing the ATP concentrations inside the granules.

### Fused in sarcoma granules diminish after prolonged coexistence of valosin-containing protein

The FUS P525L granules containing VCP disappeared in the prolonged cultured cells (Fig. 5A). The population of the cells harboring granules (granule-positive cells) among the cells overexpressing FUS P525L alone did not change in prolonged culture (40–44 h) compared to that in short-term culture (Fig. 5A). The cells overexpressing both FUS P525L and VCP reduced the population of granule-positive cells (Fig. 5A). Intriguingly, the cells expressing FUS in ATP depletion showed a smaller change in the population of the granule-positive cells after prolonged culture than the case for the cells with VCP expression (Fig. 5A). The time-dependent reduction in the population of the granule-positive cells is obvious for the cells expressing VCP.

**Figure 5:**
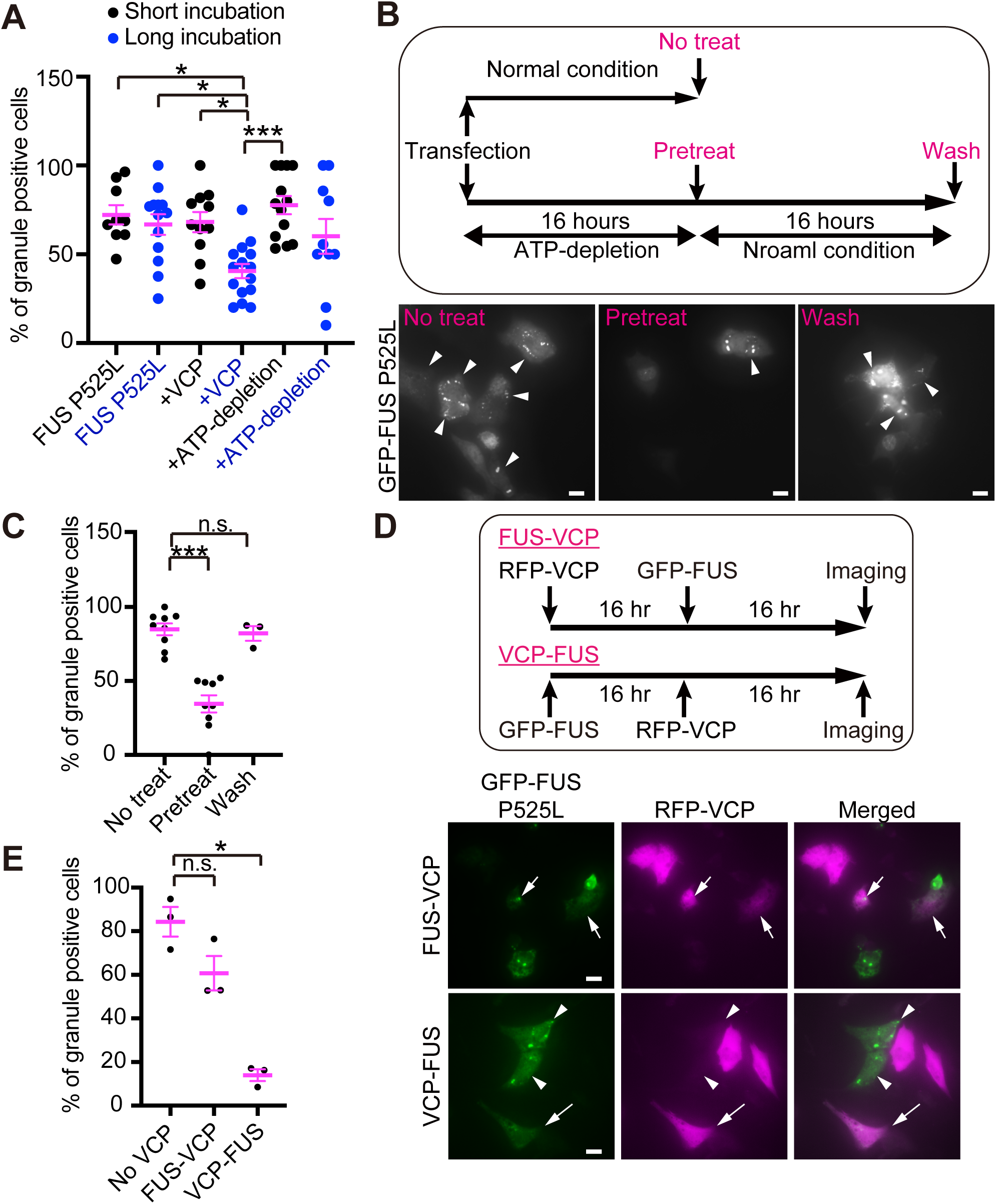
VCP amount and ATP concentration affect the formation and clearance of FUS granules. (A) The graph shows the differences in the population of granule-positive cells among cells with indicated manipulation. The percentage was calculated by counting the number of granule-positive cells over transfected cells in a field. *P < 0.05, ***P = 0.0003, and others were P > 0.35 by Dunn’s multiple comparisons test by comparing among all samples. n = 9, 13, 11, 15, 13, or 10 from left to right. Error bars show s.e.m. The data were obtained from two independent experiments. (B) (top) Scheme of the experimental procedure. (bottom) Representative images of cells observed at the timing indicated with magenta characters in the scheme. Arrowheads indicate the cells with FUS granules. Bars on images show 5 μm. Pretreat means that cells were treated with CCCP and 2DG, the ATP-depletion condition, simultaneously with transfection. Wash means that the cells were cultured with fresh media without CCCP and 2DG after observing “Pretreat”. (C) The graph shows the differences in the population of granule-positive cells. FUS P525L indicates the cells expressing GFP-FUS P525L. The percentage was calculated by counting the number of granule-positive cells over transfected cells in a field. \*\*\**P* = 0.0006 and n.s. was *P* > 0.05 by Dunn’s multiple comparisons test by comparing No treat to others. n = 9, 9, and 3 from left to right. Error bars show s.e.m. The data were obtained from two experiments. (D) (top) Scheme of the experimental procedure. FUS-VCP means that FUS overexpressed prior to VCP overexpression, and VCP-FUS means VCP overexpressed prior to FUS overexpression. (bottom) Representative images of cells observed. Arrowheads indicate the cells expressing VCP with FUS granules. Arrows indicate cells expressing VCP without FUS granules. Bars on images show 5 μm. (E) The graph shows the differences in the population of granule-positive cells observed under the experimental condition same to (E). The percentage was calculated by counting the number of granule-positive cells over transfected cells in a field. \**P* = 0.0225 and n.s. was *P* = 0.5934 by Dunn’s multiple comparisons test by comparing No VCP to others. n = 3 for all, from left to right. Error bars show s.e.m. The data were obtained from three experiments.

ATP depletion suppressed *de novo* FUS granulations (Fig. 5D). The population of the granule-positive cells diminished when cells were pretreated to be ATP depletion with CCCP (Fig. 5D). Removing CCCP restored FUS granulations (Figs. 5C and 5D). The data demonstrate FUS granulation requires a certain ATP concentration, consistent with the reported *in vitro* finding (Kang et al., 2018).

VCP expression in the cells harboring the FUS granules (Fig. 5D) had a minor impact on the population of granule-positive cells (Fig. 5E). Conversely, inducing FUS expression in the cells overexpressing VCP (Fig. 5D) diminished granulation (Fig. 5E). These results are consistent with the finding that cells in ATP depletion reduced FUS granulation (Fig. 5C). FUS granulation starts at 0.2 mM ATP concentrations *in vitro* (Kang et al., 2018). ATP consumption by VCP in cells reduced cytosolic ATP concentrations to the level suppressing FUS granulation.

### Fused in sarcoma granule size changes according to the intracellular adenosine triphosphate concentrations

FUS granules became smaller at lower ATP concentrations in cells overexpressing VCP or in ATP depletion (Figs. 6A and 6B). Proteolysis blocking by EerI or MG-132 showed less significant change in FUS granule size (Figs. 6A and 6B). VCP A232E with three-times elevated ATPase activity reduced FUS granule size as in the cells in ATP depletion (Fig. 6B).

**Figure 6:**
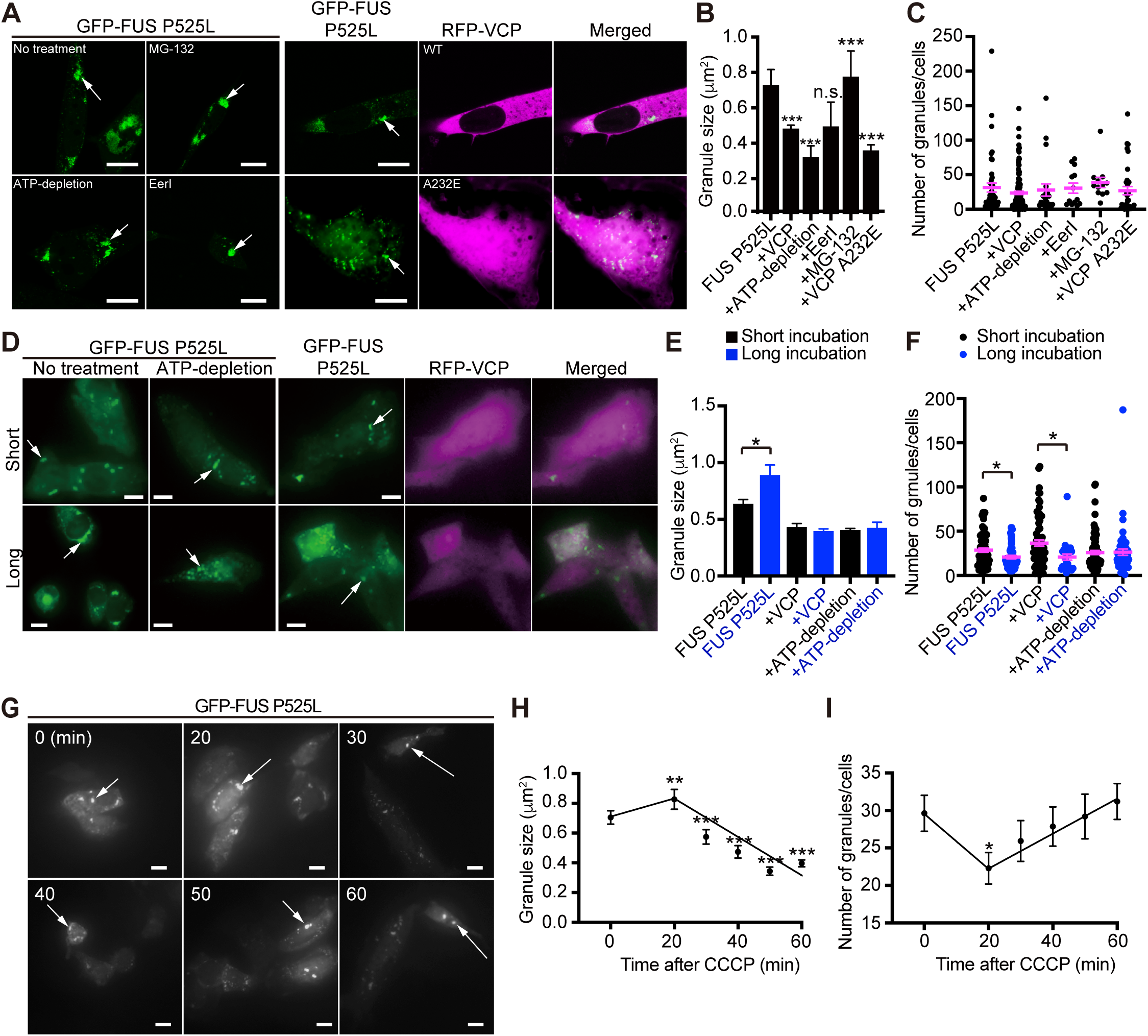
Overexpression of VCP and ATP depletion affect the morphological aspects of FUS granules. (A) Representative images of cells overexpressing GFP-FUS P525L (left) or cells also expressing RFP-VCP (wild-type, top-right group; A232E mutant, bottom-right group). The cells overexpressing only GFP-FUS P525L were also treated as indicated within the panel. (B) The size of the mutant FUS granules with indicated manipulation. \*\*\**P* < 0.0001 or n.s. (*P* = 0.7696) by Dunn’s multiple comparisons test by comparing FUS P525L to others, mean*±*s.e.m., n = 1385, 2875, 2081, 659, 311, and 946 from left to right. (C) The number of mutant FUS granules with indicated manipulation. Each plot indicates the number of granules in one cell. *P* > 0.05 by Dunn’s multiple comparison test by comparing FUS P525L to others, mean±s.e.m., n = 44, 126, 21, 13, 12, and 35 from left to right. The data were acquired from two independent experiments in samples of +ATP-depletion, +EerI, +MG-132, and +VCP A232E and three independent experiments in the samples of FUS P525L and +VCP in (B) and (C). (D) Representative images of cells overexpressing GFP-FUS P525L (left four images) or cells also expressing RFP-VCP (right six images) of short-or long-term incubation. A group of cells expressing GFP-FUS P525L were also treated with CCCP and 2DG (indicated as ATP-depletion). Arrows indicate one of the FUS granules. (E) The graph shows the size of granules among cells with indicated manipulation. \**P* = 0.0377 and others were *P* > 0.05 by Mann-Whitney test between normal and longer incubation. mean*±*s.e.m., n = 421, 219, 591, 298, 324, or 119 from left to right. (F) The graph shows the number of granules among cells with indicated manipulation. \**P* = 0.0396 in FUS P525L samples, \**P* = 0.0102 in +VCP samples, and *P* > 0. 5 in +ATP-depletion samples by Mann-Whitney test between normal and longer incubation. (G) Representative images of cells expressing GFP-FUS P525L at the indicated time after being treated with CCCP and 2DG for ATP depletion. Arrows indicate one of the granules. (H) The plots of the mean granule sizes against time after ATP-depletion. The black line indicates the linear regression from time 0 to time 20 min and from time 20 min to time 60 min. **P = 0.0012 or ***P < 0.0001 by Dunn’s multiple comparisons test by comparing time 0 to other time points. n = 403 at time 0, n = 309 at time 20 min, n = 337 at time 30 min, n = 404 at time 40 min, n = 437 at time 50 min, n = 709 at time 60 min. (I) The plots of the mean granule number in one cell against time after ATP-depletion. The black line indicates the linear regression from time 0 to time 20 min and from time 20 min to time 60 min. \**P* = 0.0255 or n.s. (*P* > 0.3) by Dunn’s multiple comparisons test by comparing time 0 to other time points. n = 68 at time 0, n = 70 at time 20 min, n = 65 at time 30 min, n = 72 at time 40 min, n = 74 at time 50 min, and n = 113 at time 60 min. The data were obtained from two independent experiments in (E), (F), (H), and (I). Error bars show s.e.m. Bars on images show 10 μm.

The number of granules in the cells was less sensitive to intracellular ATP concentrations (Fig. 6C). Proteolysis blockers, EerI and MG-132, did not change the number of granules in the cells (Fig. 6C).

The prolonged-culture (40–44 h) cells enhanced the change in the number of granules (Fig. 6D). FUS granules in prolonged culture cells became larger (Fig. 6E), thus reducing the number of granules (Fig. 6F). FUS granules became fused during the extended culture. FUS granules in the cells overexpressing VCP showed a marginal change in granule size (Fig. 6E) but significantly reduced the number in the prolonged cultured cells (Fig. 6F). Continuing ATP consumption by VCP reduces ATP concentrations in cells, which probably destabilizes the granules to dissolve in the cells expressing VCP (Fig. 6F). Cells in ATP depletion showed no time-dependent change in granule size and the number of granules (Figs. 6E and 6F).

The granule size changed within 60 min after blocking ATP synthesis with CCCP (Figs. 6G and 6H). FUS granules became larger up to 20 min after the CCCP input and subsequently reduced in size (Fig. 6H). The number of granules diminished up to 20 min and subsequently increased (Fig. 6I). The changes may imply that the granules coalesced for the first 20 min after the CCCP input, then they became unstable because of the lowered ATP concentration by blocking ATP synthesis (Fig. 6H). The unstable granules break into smaller granules, thus raising the number of granules in cells (Fig. 6I).

The swift change in the FUS granule morphology in cells (Figs. 6H and 6I) demonstrates that FUS granulation is sensitive to ATP concentrations. ATP concentration in the cells treated with CCCP will gradually diminish under blocking ATP synthesis. ATP concentration for the first 20 min will promote FUS granulation. In the following time, ATP concentration will continuously diminish to destabilize the granules. It is conceivable that VCP co-localized to the granules similarly regulates the FUS granulation in a time-dependent manner by continuing intragranular ATP consumption, which should be a biphasic change as found in FUS granulation progressing in the cells under blocking the ATP synthesis (Figs. 6H and 6I).

## Discussion

This study revealed a new role of VCP, a AAA ATPase, in FUS granulation in cells. VCP in the FUS granules stabilized *de novo* granules by rigidifying the liquid-like outer sphere by reducing intragranular ATP concentrations (Fig. 7A); however, the prolonged stay of VCP in the granules destabilized the FUS granules to dissolve by continuing intragranular ATP consumption with its ATPase activity (Fig. 7B).

**Figure 7:**
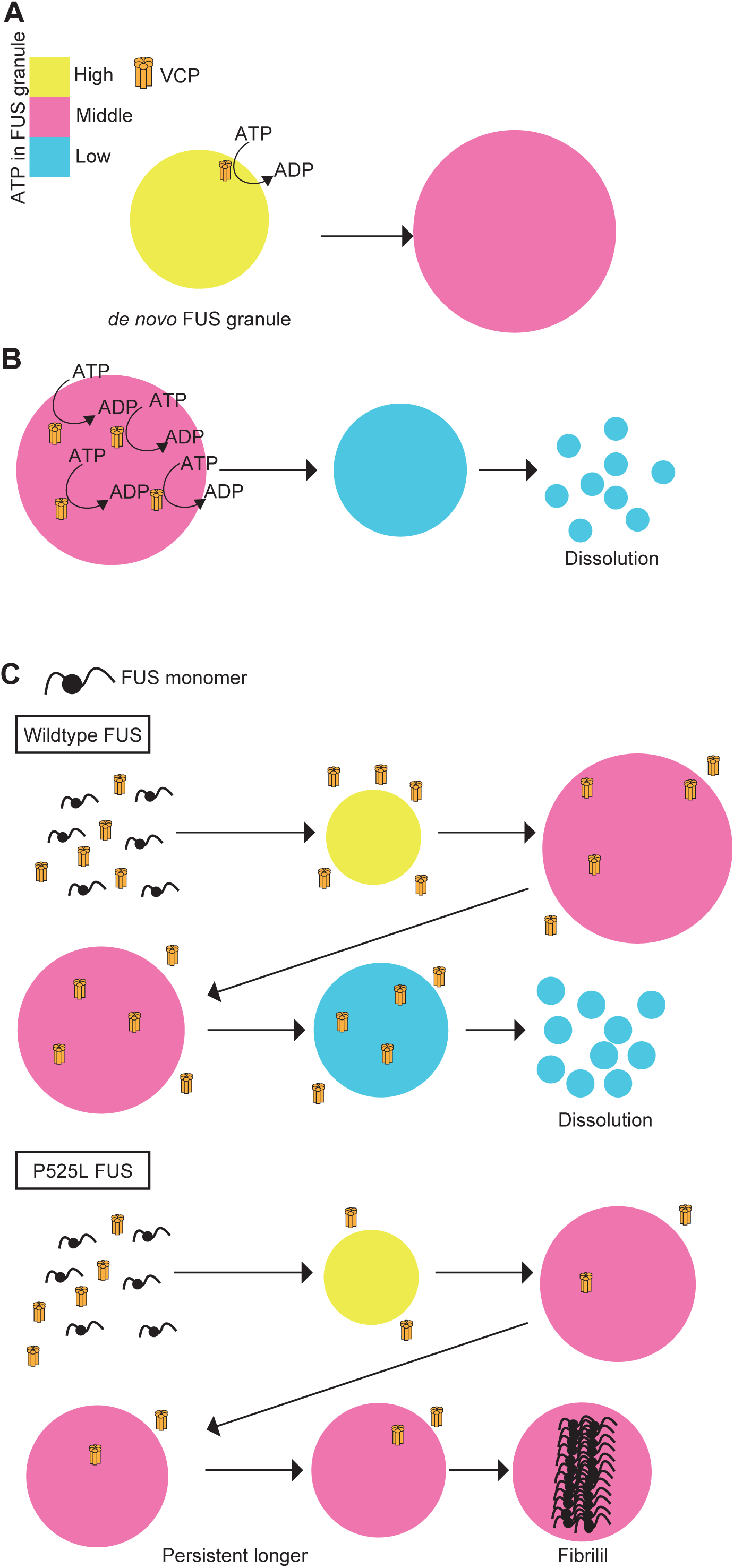
Regulatory role of VCP in FUS granulation. (A) FUS granules in the cytosol contain high ATP concentrations around 4 mM (level indicator in yellow) (Yoshida et al., 2016), *de novo* FUS granules (yellow circle). ATP at high concentrations destabilizes FUS granules (Kang et al., 2018; Patel et al., 2017). Thus, *de novo* FUS granule is fragile. VCP co-localized to the FUS granules consumes intragranular ATP with its ATPase activity to reduce the ATP concentrations to the level that stabilizes the FUS granules (around 1 mM, level indicator in red) (Kang et al., 2018): the stabilized FUS granule is drawn as a red circle. (B) VCP co-localized to the stabilized FUS granule continues consuming ATP to reach the low-level ATP concentrations (level indicator in blue) in the granule drawn as a blue circle. FUS granulation requires at least 0.2 mM ATP concentration (Kang et al., 2018). The FUS granules at low intragranular ATP concentrations will dissolve. VCP co-localized to the FUS granules stabilizes *de novo* granules but subsequently destabilizes the granules to dissolve by continuing consumption of the intragranular ATP. VCP in the FUS granules plays as a timer to limit the lifetime of the FUS granules in cells. (C) The wild-type FUS mislocalized in the cytosol forms fragile *de novo* granules (a yellow circle). Once VCP is recruited to the *de novo* granules, VCP reduces the intragranular ATP to stabilize the granules to grow (a red circle). VCP continues ATP consumption in the granules to lower the intragranular ATP concentration to destabilize the granule (a blue circle), which will dissolve. FUS P525L with impaired affinity to VCP should have a longer turnover time. FUS P525L lacks the ability in nuclear translocation (Zhang and Chook, 2012) thus accumulates in the cytosol, which prompts FUS P525L granulation over the wild-type FUS granulation. Because of the reduced affinity to VCP, FUS P525L granules recruit a smaller amount of VCP relative to the wild-type FUS granules. FUS P525L granules with less amount of VCP (a red circle) can retain the middle range of the intragranular ATP concentrations to stabilize the granules for a longer period than the case of the wild-type FUS granules. FUS fibrillization progresses in the granules in cells (Patel et al., 2015b). The prolonged residence of the granules will promote the FUS fibrillization, which may explain the elevated fibrillation property of the FUS P525L mutant, even though FUS P525L *per se* does not show any elevated aggregation property in a test tube (Nomura et al., 2014).

The functions of the VCP in the granules rely on the two-face role of ATP in FUS granulation: ATP at low concentrations ranging from 0.2 mM to 1.0 mM facilitates FUS LLPS and dissolves the granules at concentrations exceeding 4.0 mM (Kang et al., 2019a; Kang et al., 2018; Kang et al., 2019b; Patel et al., 2017). The FUS granules are most stable at an ATP concentration of around 1.0 mM (Kang et al., 2018; Patel et al., 2015a; Patel et al., 2017).

The cytosolic ATP concentrations in mammalian cells range from 3.7 to 4.1 mM (Yoshida et al., 2016), under which ATP concentrations FUS granules are too fragile to reduce the sizes (Kang et al., 2018). The liquid-like outer spheres of the granules exposed to high cytosolic ATP concentrations will dissolve (Fig. 7A) (Shiina, 2019). The hexameric VCP contains 12 ATPase domains in a molecule (Meyer et al., 2012). VCPs in the FUS granules consume ATP to reduce the intragranular ATP concentrations, thus stabilizing the outer shells of the FUS granules (Fig. 7A), which is evident from the finding that the granules containing VCP retained their sizes even in the cells permeabilized by the detergent, digitonin (Fig. 4B).

In prolonged cultured cells, VCP in the granules keeps consuming the intragranular ATP to lower the concentrations to destabilize the granules (Fig. 7B), as evidenced by the reduced number of granule-positive cells in prolonged culture (Figs. 5A and 6F). VCP functions as a timer to limit the residence time of FUS granules in cells.

Proximity labeling with Halo-tagged FUS P525L showed that the FUS P525L granules contained a lower amount of VCP than the wild-type, implying that the FUS P525L mutant has a lower affinity for VCP (Fig. S1C). The reduced amount of VCP in FUS P525L granules was approximately 20% relative to that in the wild-type (Fig. 1D), although the amount of the immunoprecipitated VCP to FUS P525L diminished by about 50% (Fig. 1B). The finding suggests that VCP engages in the FUS granules through interactions with proteins other than FUS in the granules.

Wang et al. report that FUS P525L has a higher affinity for VCP than the wild-type with mass spectrometry (Wang et al., 2015). This discrepancy is attributed to the different sample preparation used to detect VCP–FUS interactions. The proximity labeling considers the VCP in physical contact with FUS (Mishra et al., 2020). The VCP co-purified by affinity chromatography with FLAG-His double-tagged FUS contained all the proteins in FUS granules without limiting the proteins in spatial proximity to FUS (Wang et al., 2015). The discrepant finding supports that VCP utilizes the interactions with the other components other than FUS protein.

Several studies have argued that VCP functions in SG clearance: VCP knockout by siRNAs results in a defect in the clearance of SGs, and pathogenic VCP mutants lead to constitutive accumulation of SGs in cells (Buchan et al., 2013). VCP reportedly facilitates endocytosis to degrade FUS and inhibit its toxicity: inhibition of endocytosis accumulates FUS to increase its aggregation in cells (Liu et al., 2020b).

In contrast to the VCP role in clearing SGs, this study demonstrates that VCP in the FUS granules does not use proteolysis activity but relies on ATPase activity to regulate granulation. The pathogenic VCP A232E mutant accumulates constitutive SGs (Buchan et al., 2013) because it loses its function to promote proteolysis. In the FUS granules, VCP A232E destabilized the granules with its elevated ATPase activity (Niwa et al., 2012) to promote FUS granule clearance (Fig. 6B). VCP plays different roles in SGs and FUS granules.

FUS P525L is a fibrillization-prone mutant that causes juvenile FUS pathogenesis (Conte et al., 2012; Eura et al., 2019). The raised aggregation propensity of FUS P525L in cells is ascribed to its cytoplasmic mislocalization (Zhang and Chook, 2012); FUS normally localizes to the nucleus (Lee et al., 2006). FUS P525L does not show elevated aggregation property *in vitro* (Nomura et al., 2014). FUS P525L fibrillization links to its granulation in the cytoplasm.

FUS fibrillization progresses in granules (Patel et al., 2015b). Our model proposes that VCP limits the residence duration of FUS granules in cells: (Fig. 7C). This work revealed that FUS P525L impaired the affinity to VCP (Fig. 1B) and diminished the amount of VCP co-localized in the FUS P525L granules (Fig. 1C). The reduced amount of VCP in the granules should extend the residence of the granules in cells because of the reduced intragranular ATP consumption by VCP (Fig. 7C). The prolonged residence of FUS P525L granules in cells should promote FUS fibrillization in the granules (Fig. 7C).

In conclusion, we revealed that VCP plays as a timer to limit the residence of FUS granules by consuming the intragranular ATP with its ATPase activity (Figs. 7A and 7B). FUS P525L accumulated in cytosols forms the granules containing less amount of VCP should reside longer time in cells than the wild-type FUS, which explains the elevated FUS P525L fibrillization occurring inside the granules (Fig. 7C).

This work used the VCP overexpression to explore its roles in FUS granulation, which may overestimate the plays of the VCP ATPase co-localized to the FUS granules. In reality, VCP should also play a proteolysis-promoting factor that uses ATP to balance VCP ATPase activity in regulating FUS granulation and turnover. In the future analysis, we will explore the roles of endogenous VCP in regulating FUS granulation, in particular focusing on the persistency of the FUS P525L granules in cells.

Exploring the regulatory mechanism in collaboration among the associated components like VCP in the FUS granules will unveil the self-turnover system of RNP granules in cells, which should intimately link to aberrant progression of the pathogenic fibrillization in the granules.

## Materials and methods

### Plasmids and construction

FUS wild-type and P525L mutant coding sequences cloned into the pEGFP-C1 vector were used for getting the coding genes (Yasuda et al., 2013). The wild-type and P525L mutant FUS genes were amplified from the plasmids with polymerase chain reaction (PCR).

The human VCP cDNA was reverse transcribed from total RNA by using Omniscript RT kit (Qiagen Japan, Tokyo, Japan) extracted from A549 cells (RIKEN Cell Bank, Ibaraki, Japan; authenticated) and purified with a high-purity RNA extraction kit (Qiagen Japan, Tokyo, Japan).

The Halo-FLAG-FUS in mRFP-C1 vector (Addgene, Watertown, MA) was constructed using pcDNA3 (a gift from Hyun-Woo Rhee) and the human VCP gene (NM_007126.3). A232E site-directed mutation in RFP-VCP was done with a Q5 site-directed mutagenesis kit (New England Biolabs Japan, Tokyo, Japan).

The primers used above are provided in the Supplementary Information.

The ATeam1.03-nD/nA/pcDNA3 plasmid was a gift from Takeharu Nagai (Addgene, Watertown, MA, plasmid # 51958; http://n2t.net/addgene:51958; RRID: Addgene_51958) (Kotera et al., 2010).

Plasmids used for transfection were purified using the NucleoBond Extra Midi kit (Macherey Nagel, Duren, Germany). All restriction enzymes were purchased from New England Biolabs. The competent cells (DH5α) (Toyobo, Osaka, Japan) were used for plasmid construction.

### Cell culture and transfection

NIH/3T3 cells (RIKEN Cell Bank, Ibaraki, Japan; authenticated) were grown in Dulbecco’s modified Eagle’s medium (DMEM) (high-glucose with L-glutamine and phenol red) (Gibco, Waltham, MA) supplemented with 10% fetal bovine serum (FBS) (Gibco, Waltham, MA), penicillin, and streptomycin (Gibco, Waltham, MA). A549 cells, used for RNA extraction, were grown in DMEM (low-glucose with L-glutamine and phenol red) (Gibco, Waltham, MA) supplemented with 10% FBS, penicillin, and streptomycin (Gibco, Waltham, MA). NIH/3T3 cells were used in the experiments. For overexpression, 10 g/10 cm dish of plasmids was transfected using Lipofectamine 3000 (Invitrogen, Waltham, MA). Cells were cultured for 16–20 h after transfection (short culture) and 40–44 h after transfection (long culture).

The siRNA against mouse VCP (Qiagen, catalog #: SI01469020, 5’-TCCGGTGGGCTTTGAGTCAAA-3’) was transfected with RNAi Max (Invitrogen, Waltham, MA). The cells were used for the experiments at 64 h after siRNA transfection. For the control (siControl), we used AllStars Neg. Control siRNA (Qiagen Japan, Tokyo, Japan, catalog #: 10272281). The cells with the endogenous VCP knocked down were used for exploring GFP-or RFP-tagged VCP behavior. Opti-MEM (Gibco, Waltham, MA) was used to dilute all reagents for transfection.

Cells were cultured in a medium containing 10 μM EerI (Tocris Bioscience, Bristol, UK) or 10 μM MG-132 (Chemscene, Monmouth Junction, NJ) for 10–14 h to block proteasome reactions. Cells in ATP depletion were prepared by culturing the cells in a medium containing 100 μM CCCP (R&D systems, Minneapolis, MN) and 600 μM 2-deoxy-D-glucose (2DG) (Sigma-Aldrich, St. Louis, MO) for 1 h (short culture) or 24 h (long culture).

Cell stress was induced by culturing the cells in a medium containing 500 μM arsenite sodium (Merck, Darmstadt, German) for 30 min. Stress induction to the cells in ATP depletion used the cells pre-cultured in the medium containing CCCP and 2DG as described above prior to the culture in the arsenite-containing medium.

In the cell permeabilization experiments, cells were cultured in FluoroBrite DMEM (Thermo Fisher, Waltham, MA) containing 0.05% digitonin during the observation (20 min).

### Immunofluorescence (IF) and western blotting (WB)

The following primary antibodies were used: Mouse monoclonal anti-FLAG M2 (Sigma Aldrich, St. Louis, MO, catalog #: F1804, diluted to be 1:1000 for WB), rabbit polyclonal anti-FUS (Proteintech, Rosemont, IL, catalog #: 11570-1-AP, diluted to be 2.5 ng/μl for IF and 0.25 ng/μl for WB), rabbit polyclonal anti-VCP (Proteintech, Rosemont, IL, catalog #: 10736-1-AP, diluted to be 1.3 ng/μl for IF and 0.13 ng/μl for WB), mouse anti-Transportin 1 (clone #: D45, Abcam, Cambridge, UK, catalog #: ab10303, diluted to be 2.0 ng/μl), mouse monoclonal anti-GAPDH (Medical and Biological Laboratories, Aichi, Japan, catalog #: M171-3, diluted to be 0.15 μg/μl for WB), mouse anti-G3BP (Abcam, Cambridge, UK, catalog #: ab56574, diluted to be 2.5 ng/μl for IF), and mouse Ubiquitin antibody conjugated with horseradish peroxidase (HRP) (clone #: P4D1, Cell signaling technology, Danvers, MA, catalog #: 14049, diluted to be 1:1000). We used the following secondary antibodies: anti-mouse IgG conjugated with HRP (Proteintech, Rosemont, IL, catalog #: SA00001-12, diluted to 1:5000), anti-rabbit IgG conjugated with HRP (Invitrogen, Waltham, MA, catalog #: 31460, diluted to be 0.16 ng/μl), anti-rabbit IgG conjugated with Alexa 488 (Invitrogen, Waltham, MA, catalog #: A-21206, diluted to 5 ng/μl), and anti-mouse IgG conjugated with Alexa 647 (Invitrogen, Waltham, MA, catalog #: A-21236, diluted to 5 ng/μl).

In the IF experiments, the cells were chemically fixed in 4% formaldehyde in phosphate-buffered saline (PBS) for 15 min, permeabilized in 0.2% Triton X-100 in PBS for 5 min, and blocked in 5% FBS in PBS for 60 min. Primary antibodies diluted in blocking solution were incubated with the samples overnight at 4 °C. The secondary antibody labeled with Alexa Fluor 488 or Alexa Fluor 647 diluted in blocking solution was incubated with the cells for 1 h at room temperature (24-25°C). The samples were washed three times after each immunoreaction for 5 min. Fluoromount-G mounting medium with 4′,6-diamidino-2-phenylindole (Invitrogen, Waltham, MA) was used to mount the samples.

The cell lysate was prepared using NIH/3T3 cells cultured in a 10 cm dish at 80% confluence. The cells were washed twice with PBS and harvested in 300 μl lysis buffer (50 mM Tris-HCl, pH 7.4, 1% (w/v) Triton X-100, 0.5%(w/v) sodium deoxycholate, 150 mM NaCl, 0.1% (w/v) sodium dodecyl sulfate [SDS]) supplemented with a protease inhibitor cocktail (Nacalai Tesque, Kyoto, Japan). Cells were lysed in lysis buffer at 4 °C for 30 min. Cell debris, including the chromosome fraction, was removed by centrifugation (16,630 x g for 10 min). The protein concentration in the supernatant was measured using the BCA assay (Thermo Fisher, Waltham, MA). The supernatant protein concentrations were adjusted to the same level among the samples. The supernatant boiled at 98 °C for 5 min was subjected to SDS-polyacrylamide gel electrophoresis (PAGE) with 7.5% polyacrylamide gel. Protein bands were transferred to polyvinylidene difluoride membranes (Bio-Rad, Hercules, CA) and subsequently rinsed with Tris buffer Saline with treen-20 (TBST; 20 mM Tris-HCl, pH 7.4, 150 mM NaCl, 0.1% (w/v) Tween-20, and 0.5% (w/v) bovine serum albumin) for blocking. The primary antibody was reacted at 4 °C overnight, followed by secondary antibody binding at room temperature (24-25°C) for 1 h. Triplicate washing with TBST for 5 min was performed after each antibody reaction step. Target proteins were detected using Chemi-Lumi One Ultra (Nacalai Tesque, Kyoto, Japan). Stained gels were imaged with a chemiluminescence imager, Omega Lum G (Gel Company, San Francisco, CA).

The cell lysate was used for the immunoprecipitation assay. After centrifugation, 20 μl supernatant was used to monitor the total amount of target protein (input). Dynabeads conjugated with anti-FLAG antibody (Sigma Aldrich, St. Louis, MO, catalog # M8823) were used to pull down the proteins bound to the FLAG-tagged bait protein, and the anti-FLAG antibody-conjugated beads were washed twice with the lysis buffer prior to use. The supernatant was mixed with the prepared Dynabead using a rotating device for 2 h at 4 °C. A 20 μl supernatant was used for the flow-through fraction (FT) in the analysis. The collected beads by centrifugation were washed with lysis buffer four times to obtain the proteins bound to the bait.

### Photo-cross-link of Halo-FLAG-fused in sarcoma and its proximal protein (Prof. Rhee)

The photoreactive ligand used in this study is a gift from Prof. Hyun-Woo Rhee (Mishra et al., 2020). HeLa cells were transfected with Halo-FLAG-FUS plasmids using the FuGENE HD Transfection Reagent (Promega, Madison, WI, catalog #: E2311). The cells were cultured in a medium containing the ligand (10 μM) for 1 h at 37 °C. The cells were rinsed twice with PBS and subjected to UV irradiation at 20000 μJ/cm^2^ for 10 min using a UV-cross linking device (UVP, CL-1000, analytikjene, Jena, German). The cell lysate from the UV-irradiated cells was subjected to immunoprecipitation with an anti-FLAG antibody. SDS-PAGE was used to analyze the proteins collected by immunoprecipitation with Coomassie brilliant blue (CBB) staining.

A portion of the SDS-PAGE gel containing proteins exceeding 140 kDa was subjected to in-gel digestion to extract proteins that were subsequently condensed by evaporating solvent and dissolved in 0.1% formic acid in water. The protein solution was analyzed using LC-MS/MS. The MAS fingerprint analysis of the collected proteins was performed using the software (Scaffold_4.8.8, Proteome Software Inc.). The changes in the amount of protein were elucidated based on the signal intensities.

### Microscopic image analysis

Cells were cultured on a glass bottom dish (Matsunami, Osaka, Japan) and in FluoroBrite DMEM medium (Thermo Fisher, Waltham, MA) to collect live-cell images. FRAP experiments were performed using an inverted laser scanning microscope (FV1000, Olympus, Tokyo, Japan) with an incubator (CytoGROW™ G.L.P., Sanyo, Osaka, Japan) to culture the cells during observation. A 60× objective oil lens (NA: 1.49, PLAPON, Olympus, Tokyo, Japan) was used. A 41.98 × 41.98 μm^2^ window (1024 × 1024 pixels) was used for imaging. The collected images were 10× averaged in the Z-stack mode. EGFP-tagged proteins were observed using 473 nm excitation and observed with a band-pass filter (485–545 nm). RFP-tagged proteins were detected with a band-pass filter (583–683 nm) by 559 nm excitation. The similar microscope setup was used to image the chemically fixed cells.

In the experiments using ATeam, the fluorescence excited by 440 nm light was split into CFP and YFP-derived fluorescence using a dichroic mirror (DM 405/473/559/635). The fluorescence from CFP and YFP was selectively detected using a corresponding band-pass filter. RFP-FUS proteins in cells harboring ATeam were excited at 635 nm to avoid unwanted YFP excitation.

The collected images were analyzed using the Fiji software (Image-J). The background noise was estimated by averaging the signals in the area where the cells were absent for each channel. Granules were identified in the images based on the fluorescence intensities over the threshold (three times higher intensity than the background level). The selected granule images were used to count the number of granules and estimate their size distributions. The signal intensity ratio I(CFP)/I(YFP) was estimated using the program ImageJ plugin ratio plus (Morikawa et al., 2016).

### FRAP experiments

In the FRAP experiments, we collected the images in a 10.56 × 0.56 μm^2^ window (64 × 64 pixels) (temporal resolution: 0.01 s) (Morikawa et al., 2016). The photobleaching area was set to 3 × 3 μm^2^ (20 × 20 pixels) (Goehring et al., 2010; Morikawa et al., 2016). Target granules were selected by visual inspection, and their sizes were sufficient to cover more than half of the illumination spot area. After a two-second pre-bleaching, a two-second illumination was applied for photobleaching at the maximum laser power. The photorecovery process was monitored for 8 s. The averaged target area intensity during the pre-bleaching period was used as a reference for the intensities after photobleaching.

One-phase exponential fitting was applied to the intensity recovery profile to determine the half-intensity recovery time (*t*_1/2_) to elucidate the diffusion coefficient *D* according to the following relation:

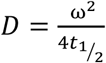

where, ω^2^ is the photobleached area.

### Statistics

Statistical analysis of the data was performed using Prism 8 (GraphPad Software). We used a non-parametric test to compare the data sets that did not pass the Shapiro-Wilk normality test. We applied the Mann-Whitney test to compare two datasets collected in a non-paired sampling manner. The Wilcoxon matched-pairs signed-rank test was applied to compare two datasets collected in paired sampling (WB data).

Dunn’s test was used to compare three or more datasets to consider the variance among them. Non-parametric Spearman correlation coefficients were used to explore the correlations among a series of data.

## Acknowledgment

We are grateful to Dr. Stavroula Mili (National Institutes of Health) for sending us the GFP-FUS plasmids.

## Author contributions

Conceptualization; KY, and S.-i.T; methodology; TMW, HWR, and KY; Technical support; MGK and JKS; investigation; KY; resources; HWR, TMW, and KY; writing-original draft; KY; writing-reviewing and editing; KY, TMW, HWR, and S-i.T; supervision; S.-i.T.; project administration; KY and S.-i.T.

## Supplementary Information

### Primers used in plasmid constructions

The wild-type and P525L mutant FUS genes were amplified by polymerase chain reaction (PCR) from the GFP-FUS plasmid gifted by Dr. Stavroula Mili (National Institute of Health, USA) (Yasuda et al., 2013). The primers used for the gene amplification with introducing EcoRV and XbaI restriction sites are shown below:

Forward primer: 5′-cccGATATCgATGGCCTCAAACGATTAT-3′

Reverse primer (wt): 5′-ccTCTAGATTAATACGGCCTCTCCCTGCGA-3′

Reverse primer (P525L): 5′-ccTCTAGATTAATACAGCCTCTCCCTGCGA-3′

The human VCP gene (NM_007126.3) was amplified from human cDNA by PCR using the following primers containing SacI or BamHI restriction sites:

Forward primer: 5′-cccGAGCTCAAATGGCTTCTGGAGCCGATTC-3′

Reverse primer: 5′-cccGGATCCTTAGCCATACAGGTCATCAT-3′

A232E site-directed mutation was applied to RFP-VCP using a Q5 site-directed mutagenesis kit (New England Biolabs Japan, Tokyo, Japan) with the following primers:

Forward primer: 5′-CTCTTTAAGGaAATTGGTGTGAAGCC-3′

Reverse primer: 5′-GGCAGGATGTCTCAGGGG-3′

## Supplementary figures

**Figure S1:**
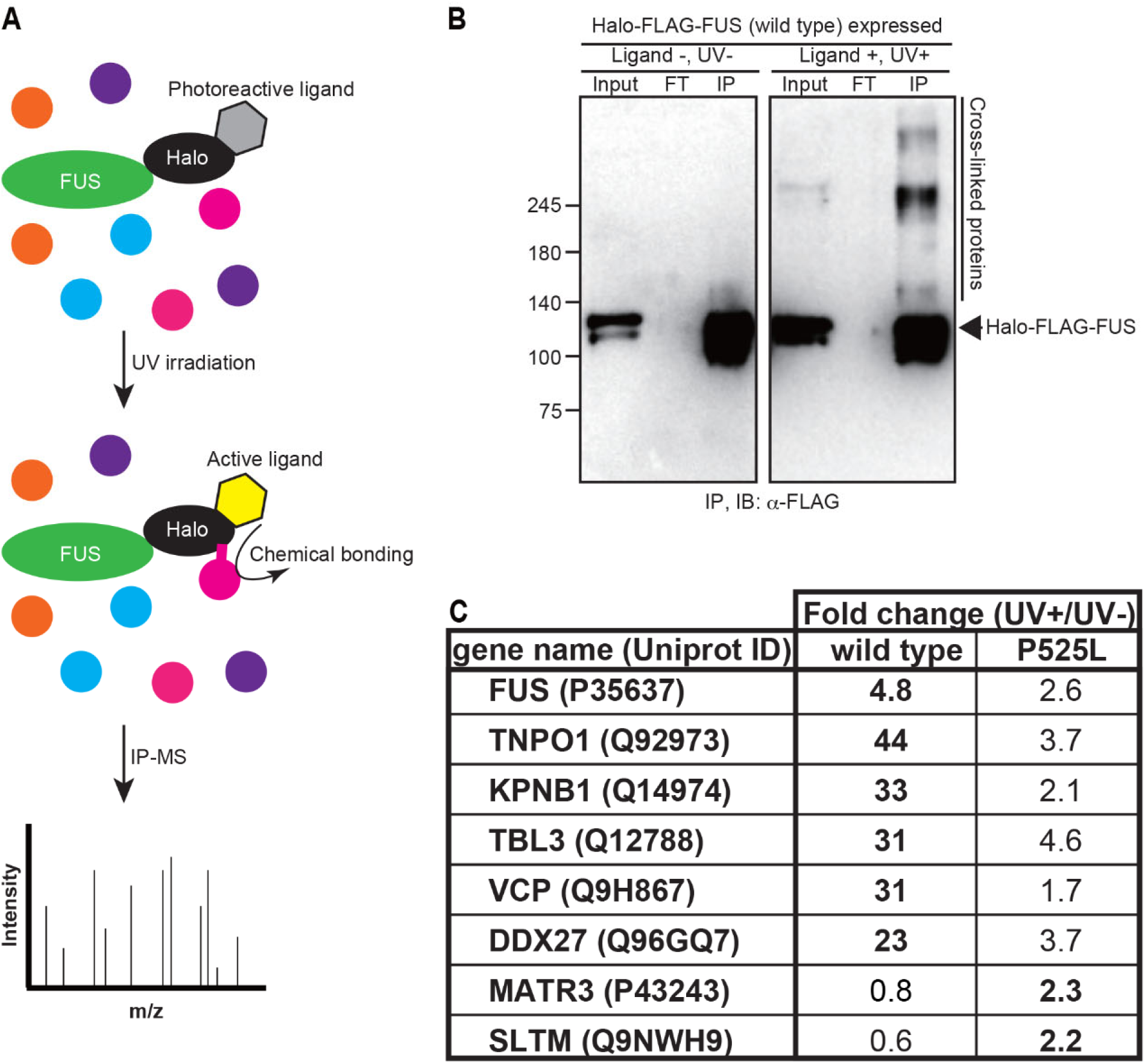
Identification of VCP as a FUS interactor which prefers to bind to wild-type FUS. (A) Scheme of photoactivatable proximity labeling (*spotlight)*. Photoreactive Halo-ligand is able to bind to Halo-tag. UV irradiation activates the ligand, and the active ligand induces the chemical bonding between Halo-FLAG-FUS protein and nascent protein. The cross-linked protein can be identified with immunoprecipitation followed by mass spectrometry. (B) Western blotting for immunoprecipitated samples of FLAG-tagged FUS protein with and without ligand-induced photo-cross-link. The lysate without precipitation (input), supernatant after precipitation (FT), or precipitates (IP) were loaded and detected with anti-FLAG antibody after immunoprecipitation with anti-FLAG antibody. The observed bands above 100 kDa are Halo-FLAG-FUS. The bands larger than Halo-FLAG-FUS size were only observed in photo-cross-linked and immunoprecipitated samples, demonstrating the cross-link-derived increase in molecular size. (C) A brief summary of mass spectrometry results. Fold change was calculated by comparing the precursor intensity of non-irradiated samples and irradiated samples. Halo-tagged wild-type FUS or FUS P525L was then compared to identify the specific binding partner to wild-type or FUS P525L protein.

**Figure S2:**
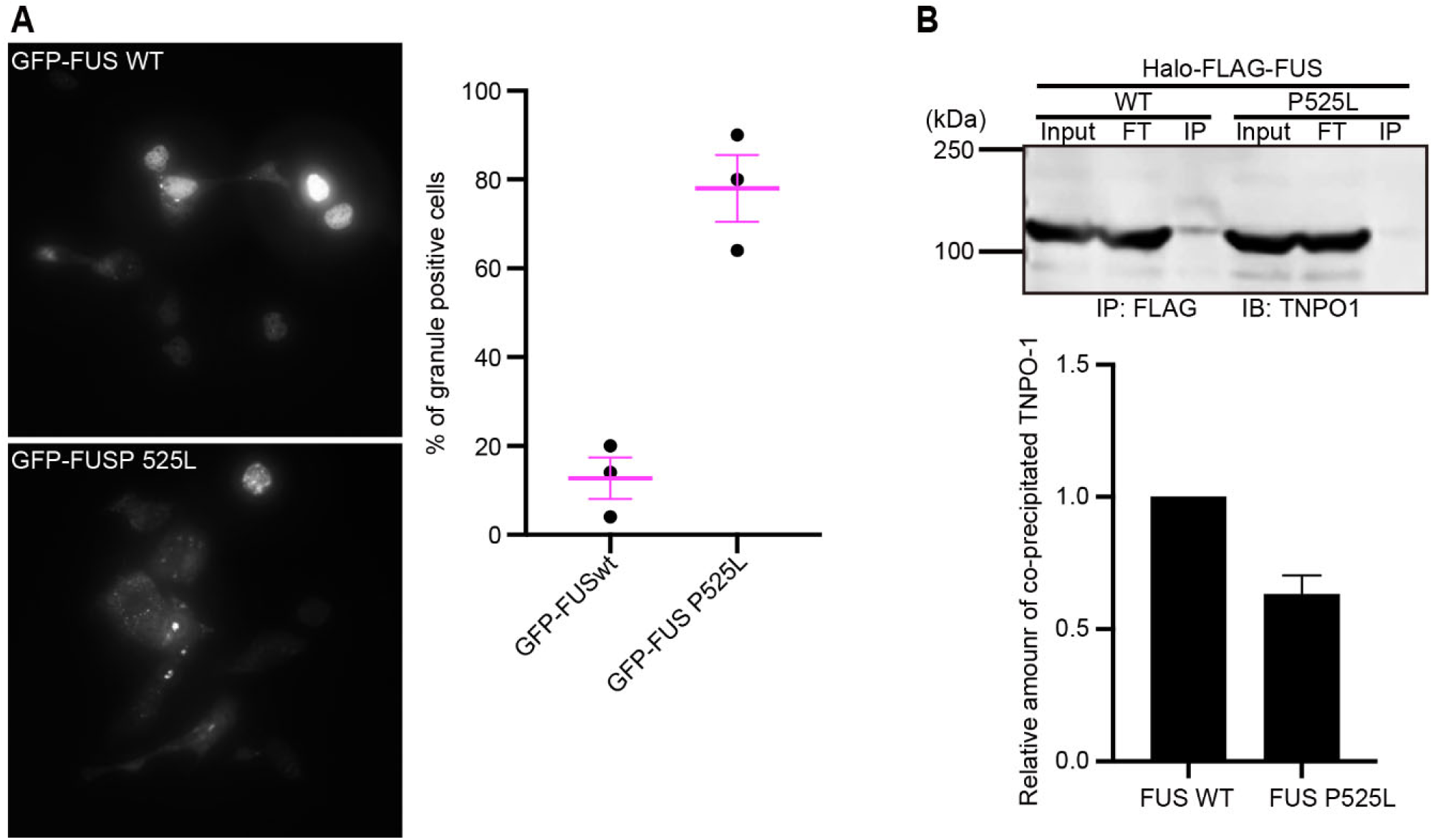
FUS granule formation is enhanced by P525L mutation on FUS, and the FUS P525L interacts less with TNPO1 than wild-type FUS. (A) Representative image of overexpressed FUS protein with (bottom) or without (top) mutation. The graph shows the average number of granule-positive cells from three trials. (B) Western blotting for immunoprecipitated samples of FLAG-tagged FUS protein overexpressed in cells. The lysates used in the experiment in Figure 1A were loaded and detected with the anti-TNPO-1 antibody. The observed bands above 100 kDa are TNPO-1, which was detected less with mutant FUS than with wild-type FUS in immunoprecipitation fraction. The graph shows the relative amount of TNPO-1 precipitated with Halo-FLAG-FUS. The intensities of TNPO-1 bands were divided by the precipitated Halo-FLAG-FUS intensity. This experiment was performed twice with different lysates.

**Figure S3:**
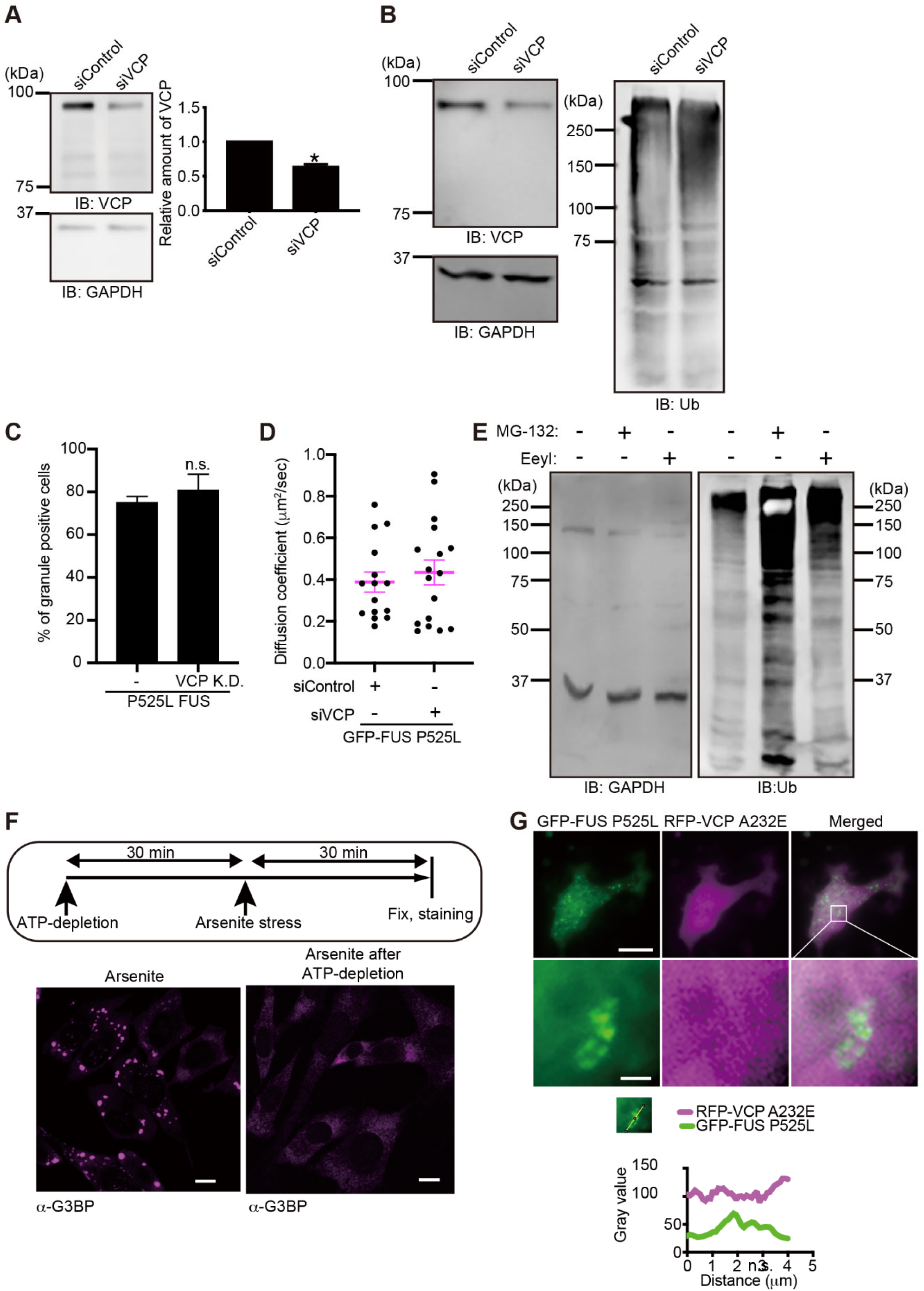
Manipulation of VCP expression, proteasome pathway, or intracellular ATP concentration. (A) Western blotting to detect VCP in cells with or without VCP knockdown (left, top), and the graph shows the differences among the amounts measured using the intensity of bands (right). The intensity of detected VCP was normalized with the intensity of detected GAPDH as the loading control (left, bottom) and sequentially normalized with the normalized VCP intensity of control samples transfected with nothing. \**P* = 0.0312 by Wilcoxon matched-pairs signed test from six independent blots. (B) Western blotting to detect VCP in cells with or without VCP knockdown (left, top), the GAPDH as the loading control (left, bottom), and the ubiquitinated proteins (right). The cells with VCP knockdown showed more poly-ubiquitinated protein signals than the control cells. (C) The population of granule-positive cells of indicated samples was calculated as the percentage of granule-positive cells over total transfected cells. No significant difference was detected in Dunn’s multiple comparison test among all samples (P > 0.9999), n = 3 for all from independent trials. (D) The diffusion coefficient of FUS granules with or without VCP knockdown. No significant difference was detected by Dunn’s multiple comparison test by comparing siControl to siVCP of FUS P525L granule (*P* = 0.1400). mean±s.e.m., n = 15 for siControl and 17 for siVCP, from two independent experiments. (E) Western blotting of cells treated with MG-132 or EerI. GAPDH (left) was used as the loading control, and ubiquitinated proteins were detected to see the inhibitory function of MG-132 or EerI. Both treatments increased the content of poly-ubiquitinated proteins. (F) Scheme of ATP-depletion upon stress induction (top) and the immunostaining of G3BP and VCP to confirm the formation of SGs. ATP-depletion completely disrupted the formation of SGs. (G) Representative images of VCP A232E-expressing cells and the plot of intensity across the granule. Whole-cell images (top) and magnified images of the granule (bottom) are shown. A plot of intensity was generated along the granule indicated as a small image above the graph. VCP signals (magenta) did not resemble the peaks of the FUS granule (green).

**Figure S4:**
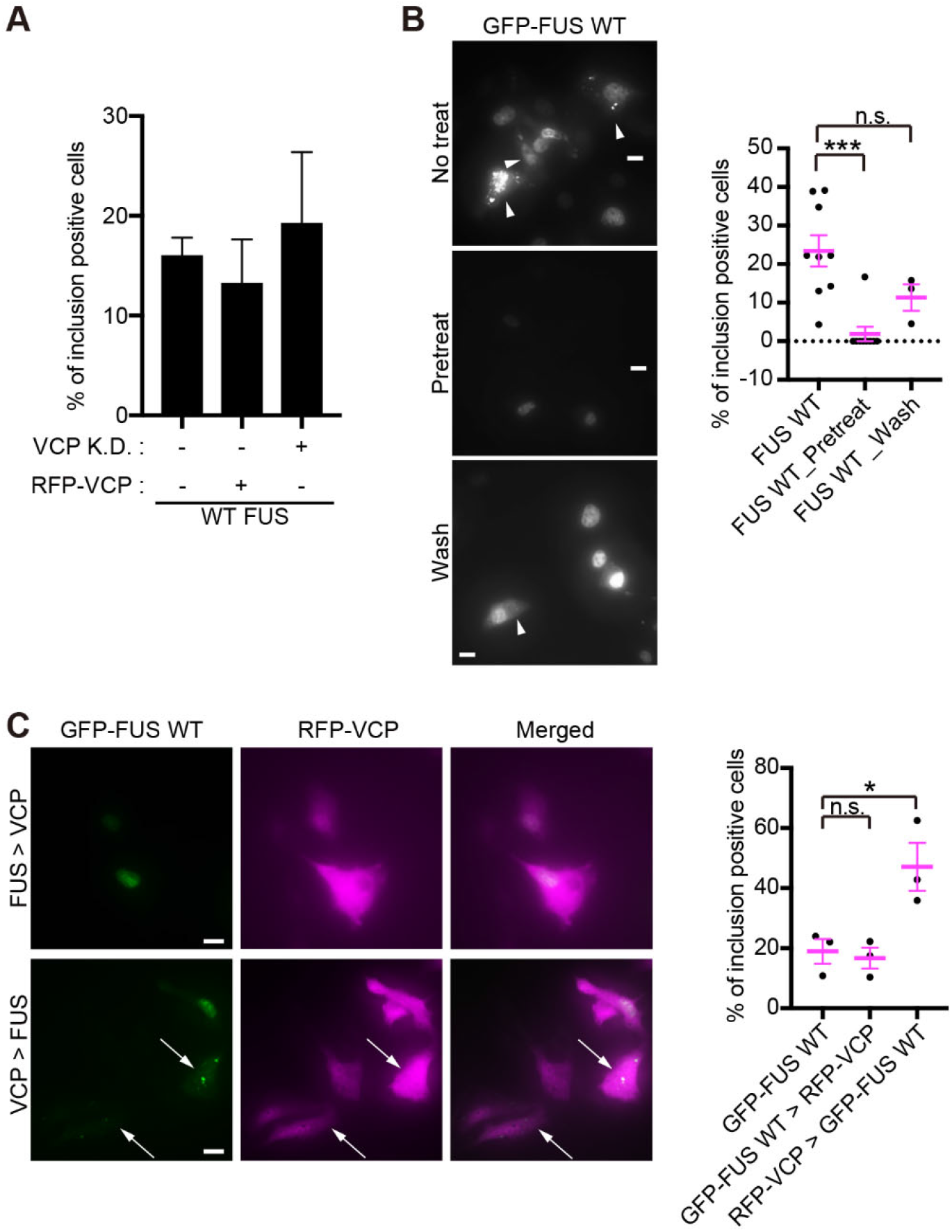
Effect of VCP on wild-type FUS granule formation. (A) (left) Representative images of cells observed. FUS-VCP means FUS overexpressed prior to VCP overexpression. VCP-FUS means VCP overexpressed prior to FUS overexpression. Arrows indicate the cells expressing VCP with FUS granules. Bars on images show 5 μm (right). The graph shows the differences in the population of granule-positive cells. The percentage was calculated by counting the number of granule-positive cells over transfected cells in a field. *P* > 0.05 by Dunn’s multiple comparisons test by comparing No VCP to others. n = 3 for all. Error bars show s.e.m. The data were obtained from three experiments.

